# GsMTx-4 Reduces Mechanosensitivity in a Model of Schwannomatosis-related Pain

**DOI:** 10.1101/2025.02.09.637313

**Authors:** Carson Gutierrez, Randy Rubright, Kimberly Laskie Ostrow

## Abstract

Patients with schwannomatosis (SWN) develop multiple tumors along major peripheral nerves, with most experiencing significant pain, though each patient’s symptoms are unique. Neuropathic, nociceptive, and inflammatory pain types have been reported, but many patients describe severe pain when a schwannoma is palpated or even lightly touched. Currently, the only effective treatment for pain relief is surgical removal. We are investigating the root causes of tumor-induced pain. In some cases, tumor growth increases pressure on nearby nerves, resulting in pain. Additionally, schwannoma cells in culture secrete proinflammatory cytokines into the surrounding medium. This conditioned medium (CM) sensitizes sensory neurons to painful stimuli both in vitro and in vivo. When injected into the glabrous skin of a mouse hindpaw, CM from painful schwannomas increases neuron sensitivity to light touch, as demonstrated by a fourfold reduction in paw withdrawal threshold (measured using the Von Frey assay) one hour post-injection (p = 0.006), with effects persisting for 24 hours (p = 0.002).We hypothesize that this increase in sensitivity is linked to mechanosensitive ion channels (MSCs), which detect pressure and stretch. These channels can be blocked by the peptide GsMTx-4. This peptide penetrates deeper into cell membranes under mechanical pressure to block MSCs from opening without affecting other ion channels. When co-injected with CM into the mouse hindpaw, 10 µM GsMTx-4 prevents heightened sensitivity to light touch. Moreover, GsMTx-4 can reverse hyperalgesia, restoring withdrawal thresholds to baseline levels. Thus, local injection of GsMTx-4 near painful tumors presents a promising, minimally invasive therapeutic approach for SWN patients.

**Significance:** Pain is a confounding comorbidity in the multiple tumor syndrome schwannomatosis. Patients harbor benign peripheral nerve sheath tumors that rarely become malignant or cause neurological deficits. Yet, patients undergo numerous surgeries for the removal of painful tumors. A non-invasive treatment for tumor-related pain is in dire need. We are examining the small peptide GsMTx-4, a blocker of mechanosensitive ion channels, as a potential therapy for painful tumors in the context of schwannomatosis.

## Introduction

Schwannomatosis (SWN) is a syndrome in which patients develop multiple tumors along their peripheral nerves [1-3]. These tumors called schwannomas are made up of Schwann cells that encase and protect axons. Although schwannomas are typically benign and rarely become malignant the primary clinical manifestation is pain. [3, 4]. In fact most patients develop pain prior to the discovery of a tumor [3]. And the intensity of pain does not always correlate with the tumor’s the size or location [5]. Patients often try multiple pain medications, sometimes up to five different types, with limited success [6]. At present the only treatment option that is effective for pain relief is surgical resection. However, medical teams face challenges in determining which tumor is causing the pain and deciding on the appropriate number of surgeries for a patient.

We do not fully understand the etiology of SWN-related pain. Neuropathic, nociceptive, and inflammatory pain have all been described by patients [7]. Palpation or incidental touch of tumors can induce intense pain [8]. We have explored the hypothesis that mechanically induced pain is a significant feature of schwannomatosis. We speculate that pressure from schwannomas on nerves could activate mechanosensitive ion channels, leading to heightened sensitivity to light touch. Furthermore, chronic inflammation has been implicated as a potential contributor to mechanical pain [9].

Studies have highlighted the role of mechanosensitive ion channels (MSCs), in the pathophysiology of various diseases [10-20]. Specifically, MSCs are targeted by GsMTx-4, a small cysteine knot peptide derived from the venom of the *Grammostola spatulata* spider and identified in Dr. Frederick Sachs’ laboratory (SUNY Buffalo) [22]. The structural and functional properties of GsMTx-4 are well detailed in many publications from the Sachs’ lab [21-27]. Moreover, the chiral form D-GsMTx-4 effectively inhibits mechanosensitive ion channels, including Piezo1 [24, 25].

Piezo1 is expressed in sensory neurons [9, 26, 27]. Its activity can be increased by Yoda1, an agonist that elevates intracellular calcium levels [28]. Yoda1 serves as a valuable tool for studying Piezo1 function in the absence of mechanical stress. GsMTx-4 has been shown to block Yoda1-induced Piezo1 activity both in vitro and in vivo [19, 26, 27]. The unique mechanism of GsMTx-4 involves interaction with the cell membrane rather than direct binding to the ion channel. Under relaxed conditions, GsMTx-4 superficially associates with the membrane’s outer lipid monolayer. When the membrane experiences tension, the peptide embeds deeper into the bilayer, physically preventing the opening of mechanosensitive channels [29, 30]. GsMTx-4 tissue accumulation and clearance studies were performed in a Duchenne muscular dystrophy mouse model D2.mdx [28]. It is non-toxic, non-immunogenic, stable in a biological environment, and has a long pharmacokinetic lifetime [28]. The rigid scaffolding of its conformation fixes relevant side chains in space and makes it resistant to enzymatic digestion [21, 29] making it a desirable candidate for drug development.

Our lab has been focusing on tumor/nerve interactions, including examining the effects of schwannoma secreted proteins on hypersensitizing sensory neurons [7, 30]. Human schwannoma tumors were collected immediately post-surgical excision and dissociated into cell cultures. Conditioned medium (CM) from cultured cells contains secreted proteins, lipids, and extracellular vesicles. We found that conditioned media from painful schwannoma tumors contains elevated levels of pro-inflammatory cytokines, and sensitizes sensory neurons to painful stimuli [31]. Injecting this conditioned medium into the footpads of healthy C57Black mice elicited pain behaviors, including heightened sensitivity to light touch (0.02 g) [36]. In this study, we further explore the effects of painful CM on mechanosensitivity using in vitro and in vivo models of SWN induced pain, proposing that GsMTx-4 could serve as an effective inhibitor of mechanically induced pain in disease.

## Materials and Methods

### Tumor cells and conditioned media and reagents

Human schwannomas were collected from surgical cases occurring at Johns Hopkins School of Medicine, from January 2014 through March 2021. Informed, written consent was obtained prior to surgery. The Institutional Review Board (IRB) of the Johns Hopkins School of Medicine approved consent forms and study design. All research was performed in accordance with relevant guidelines/regulations and approved by the IRB Study Number NA_00069904. The diagnosis of SWN was confirmed by Johns Hopkins Pathology. Human Schwannomatosis cell lines were established from tumors as described in Ostrow et al 2015. Cell cultures were maintained in DMEM, 10% FBS, 5% penicillin/streptomycin & 2uM Forskolin). Once cells reached 80% confluence in a T-25 flask, conditioned Media was collected 48h intervals. Conditioned medium was centrifuged to remove cell debris and passed through a 0.22 micron filter and stored at -80C for future use.

GsMTx-4 was purchased from Tocris Cat #4912. Yoda-1 was purchased from Tocris Cat #5586.

### Mouse cohorts

Male 8 week old C57Black mice, purchased from Jackson labs, were used in this study. All mouse experimental procedures were approved by the Institutional Animal Care and Use Committee of Johns Hopkins University School of Medicine (Approval ID MO23M18) and followed the guidelines provided by the National Institute of Health and the International Association for the Study of Pain. Animal experiments presented in this study are reported in accordance with ARRIVE guidelines.

### Dissociated Mouse DRG cell cultures

Dorsal root ganglia (DRG) were harvested from 8 week old C57BL6J mice into Complete Saline Solution (CSS; NaCl 137mM, KCl 5.3mM, MgCl_2_-6H_2_O 1mM, Sorbitol 25mM, HEPES 10mM, CaCl_2_-2H_2_O 3mM).

DRG were dissociated with TM Liberase (Trituration Moderate; 0.35U/mL in CSS, 50mM EDTA) at 37°C for 20 minutes in a rotating wheel hybridization oven, followed by TL Liberase (Trituration Low; 0.25U/mL in CSS, 50mM EDTA, 30U/mL Papain) at 37°C for 15 minutes with rotation. Cells were centrifuged at 800g for 2 minutes, the supernatant removed, and resuspended in complete DRG medium (DMEM/F12, 10%FBS, 1% glutamine, 5% penicillin/streptomycin) containing BSA (1.5mg/mL) and Trypsin Inhibitor (1.5mg/mL). The resulting suspension was enriched for neurons by passage through a 70um mesh filter. The resulting dissociated DRG cells were then spotted in 20uL TI/BSA/DMEM on poly-L-lysine/laminin coated coverslips and left to adhere for 1hour at 37 C in an incubator. Wells containing coverslips were then flooded with complete DRG medium.

### Calcium imaging

Twenty-four hours following DRG culture establishment, culture medium was removed and replaced with SWN CM or control media incubated for an additional 48 hours at 37 °C. Three separate DRG cultures were prepared for each CM to be tested. Four coverslips containing at least 30 neurons/slip were treated with CM. A minimum of 100 DRG neurons were tested per condition. Cells were treated with CM for 48 hours before loading with 2 μM fura-2 acetoxymethyl ester (Molecular Probes) in calcium imaging buffer (CIB, containing in mM: 130 NaCl, 3 KCl, 2.5 CaCl_2_, 0,6 MgCl_2_, 10 HEPES, 1.2 NaHCO_3_, 10 glucose, pH 7.45, 290 mOsm adjusted with mannitol). Coverslips with fura-2-loaded cells were mounted on an inverted fluorescence microscope (TE200, Nikon). Images were acquired with a cMOS camera (NEO, Andor) using an excitation filter wheel (Ludl) equipped with 340 and 380 nm filters. Data were acquired using NIS Elements imaging software (Nikon).

Fluorescence changes are expressed as the ratio of fluorescence emission at 520 nm upon stimulation at 340 to that upon stimulation at 380 nm (F340/F380). Yoda-1 experiments: 1 uM Yoda-1 was dissolved in CIB. During continual imaging at ∼2 sec intervals cells were perfused with 1 uM Yoda-1 with a recovery period of washing with unmodified CIB. To examine the effects of GsMtx-4, cells were incubated with SWN CM + 5uM GsMtx-4 for 48 hours prior to Yoda-1 exposure. The datapoints were corrected for differing baseline readings of Fura-2. Perfusion with KCl at a concentration of 50mM was used to identify viable neurons. Area under the curve was calculated for each treatment.

### Footpad injections

Eight week old C57BL6J mice were placed in an immobilization cone and injected with 10uL of control media 5 uM Yoda-1, or painful SWN CM in the right hind paw using a 20g insulin syringe. For the pain prevention assay: 10 uM Gsmtx-4 mixed with control media, 5uM Yoda-1, or painful SWN CM into the left paw of the mouse. Each mouse therefore served as its own internal control for withdrawal threshold change. For the pain rescue assay, a separate cohort of mice were injected with 10uL of control media 5 uM Yoda-1, or painful SWN CM in the right hind paw using a 20g insulin syringe. Twenty fours post injection 10 uM GsMTx-4 was administered to that same paw. Ten mice were injected per conditioned media. Mice were injected in groups of 20, (2 conditions; 10 mice per condition) to simplify handling and subsequent testing. All samples were deidentified and were the same in volume, color, and viscosity. The experimenter was blinded to the code for the injections and subsequent behavioral assays to eliminate bias. Mouse numbers are based on the following consideration: 10 mice per group for behavioral comparisons (82% power to detect an average 50% difference in pain behaviors, alpha=0.05, significance level of <0.05, 2-sided). Sample size was determined using GPower software.

### Radiant Paw Heating (Hargreaves Test)

Following footpad injection, an awake mouse is acclimated 30 min within a ventilated plexiglass box on a glass surface. A radiant heat source is directed at the plantar surface of the hindpaw until the mouse withdraws its paw from the source or until cutoff is reached (defined as 2.5 times the mean response latency for wild-type mice, which is usually 5-15 sec), whichever comes first. The maximum skin temperature is approximately 55°C. Each paw is tested 5 times and averaged per mouse.

### Von Frey (VF) Assessment

We employed the simplified up/down method (SUDO) to test for hypersensitivity to light touch in the injected mice, using Von Frey filaments ranging from 0.02 g-1.4g. This method identifies the minimal force that elicits a consistent response, indicating the mechanical pain threshold. Briefly, testing is initiated at 0.6g (a mid-range force). The filament is gently pushed against the surface of the glabrous skin from below for 2 seconds. A positive response is a flinch of the leg, indicating the mouse is reacting to the stimulus. If a positive response is achieved at 0.6 g, the next lowest force is tested until the animal does not respond. If a negative response to 0.6g was recorded the next highest filament is tested. To determine paw withdrawal threshold (PWT) we employed the simple up/down reader software [32]. Mice were placed into plexiglass cages in groups of 5 on top of a wire-mesh platform and given an acclimation period of twenty minutes prior to VF. Each mouse was subjected to VF prior to injection to determine a baseline or pre-injection withdrawal threshold. Following injection, VF was performed at time intervals of 1hr or 24 hours.

### Data Analysis and Statistics

#### For calcium imaging

The area under the curve of fura-2 ratio, with subtraction of the baseline obtained prior to each stimulus, was used to compare the effects of painful vs. non-painful CM, control media (+/-Gsmtx-4). Brown-Forsythe and Welch ANOVA tests with Dunnett’s T3 multiple comparisons test were used to compare the effects of CM groups (Graphpad Prism10). An individual cell’s response to stimuli was considered to be positive if the fura-2 ratio was greater than 0.1 after baseline subtraction. p value <0.05 was considered significant.

#### For Von Frey analysis

Two-way ANOVA with repeated measures, and Tukey’s multiple comparison tests were performed to determine significance between groups using Graphpad Prism 10 for timecourse experiments. Brown-Forsythe and Welch ANOVA tests with Dunnett’s T3 multiple comparisons test were used to compare the effects of CM groups (Graphpad Prism10) for single timepoint tests.

## Results

### GsMTx-4 inhibits mechanosensitive currents in dissociated sensory neurons

We confirmed the presence of Piezo-1 in mouse Dorsal Root Ganglia neurons by RT-PCR. Piezo-1 is expressed in intact L4/L5 DRG neurons and in dissociated DRG neuronal cultures. In addition, the Piezo-1 agonist Yoda-1 does not increase expression of Piezo-1 at the mRNA level **(Figure 1)**. As a proof of concept, we examined whether painful SWN CM can induce Piezo-1 currents in dissociated DRG cells. Mouse dorsal root ganglia (DRGs) incubated with Painful CM for 48 hours. Following incubation, the neurons were subjected to fluorescent calcium imaging analysis during challenge with low-dose Yoda-1 (1uM). Yoda-1 is a chemical compound acts as a gating modifier to promote Piezo-1 calcium currents in the absence of mechanical stress [10]. Low-dose Yoda-1 evoked an increase in calcium in a subset of neurons that continued after washout and did not return to baseline levels curve **(Figure 2a)**. Area under the curve (AUC) was calculated per treatment to measure magnitude and duration of response. Painful CM sensitized DRG neurons to low dose Yoda-1 compared to CM isolated from non-painful tumors or control media as demonstrated by an increased AUC (Painful CM AUC=14, control CM AUC=8 ; p=0.01) **(Figures 2a&b)**. When 5uM GsMTx-4 was added to the cells along with painful CM, hypersensitization to Yoda-1 was significantly decreased (AUC=6; p<0.0001) to that of control CM **(Figure 2b)**, therefore demonstrating inhibition of Piezo-1 currents. To confirm its specificity, we evaluated whether GsMTx-4 could block TRPV1 activation triggered by painful CM. Our results showed that GsMTx-4 had no impact on TRPV1 activation **(Figure 3a)**, highlighting its selective inhibition of mechanosensitive channels.

**Figure 1:**
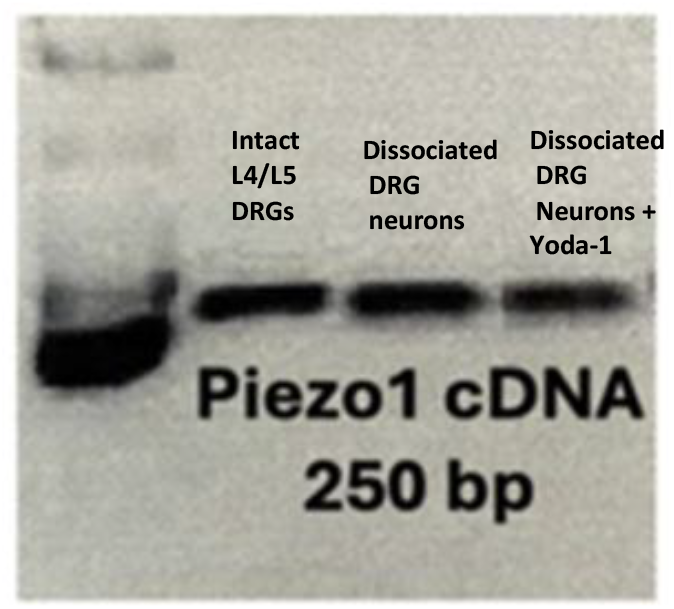
Piezo1 expression in DRG neurons. Mouse Dorsal Root ganglia were extracted from 10 mice. Expression of Piezo-1 was determined by RT-PCR. A 250 bp band corresponds to the amplified portion of Piezo-1. Expression is similar in L4/L5 intact DRGs (left lane), dissociated DRG neuronal cultures (middle lane), and in dissociated neuronal cultures treated with Yoda-1(right lane).

**Figure 2:**
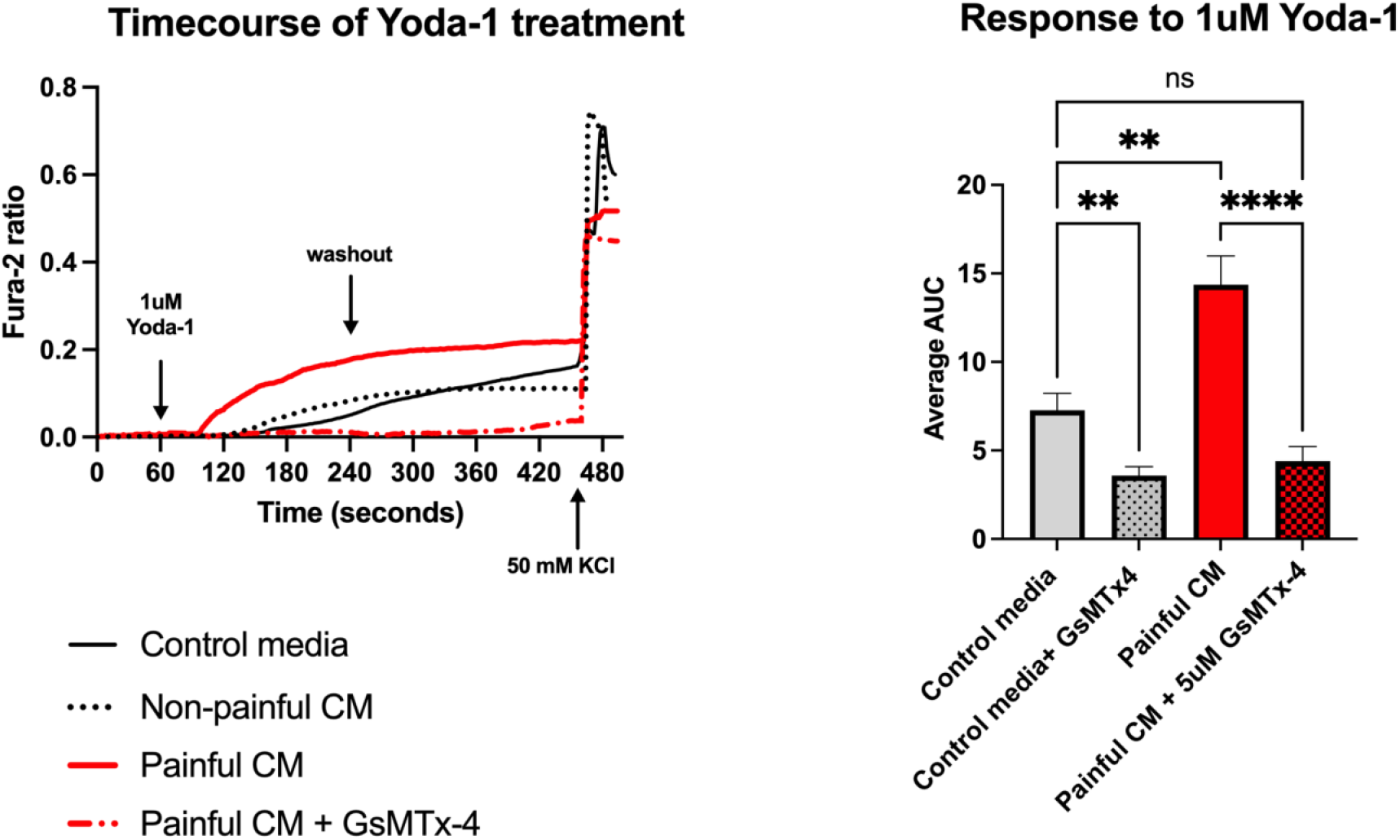
Effects of GsMTx-4 on Piezo-1 Calcium Currents. Dissociated DRG cultures were pre-treated with painful schwannoma CM +/-5uM GsmTx-4, non-painful schwannoma CM, or control media for 48 hours before imaging. Fura-2 ratio measurements were recorded at 2 second intervals. The datapoints were corrected for differing baseline readings of Fura-2. Area under the curve was calculated for each treatment. Two coverslips of DRGs (∼200 cells) were tested per condition. *Brown-Forsythe ANOVA test* with Dunnett’s T3 multiple comparisons was employed to determine statistical significance (p<0.05 is considered significant) a). Conditioned media (red solid line) from painful SWN tumors sensitize DRG neurons to the Piezo-1 agonist Yoda-1 in vitro. Non-painful tumor CM (dotted black line) had no effect on neuronal sensitivity. GsMTx-4 mixed with Painful CM inhibited Yoda-1 induced Piezo currents. **b)** Co-incubation of control media and 5 uM GsMTx-4 or painful CM and 5uM GsMtx-4 for 48 hours, decreased hypersensitivity to Yoda-1 (p<0.0001) as demonstrated by a decrease in area under the curve.

**Figure 3:**
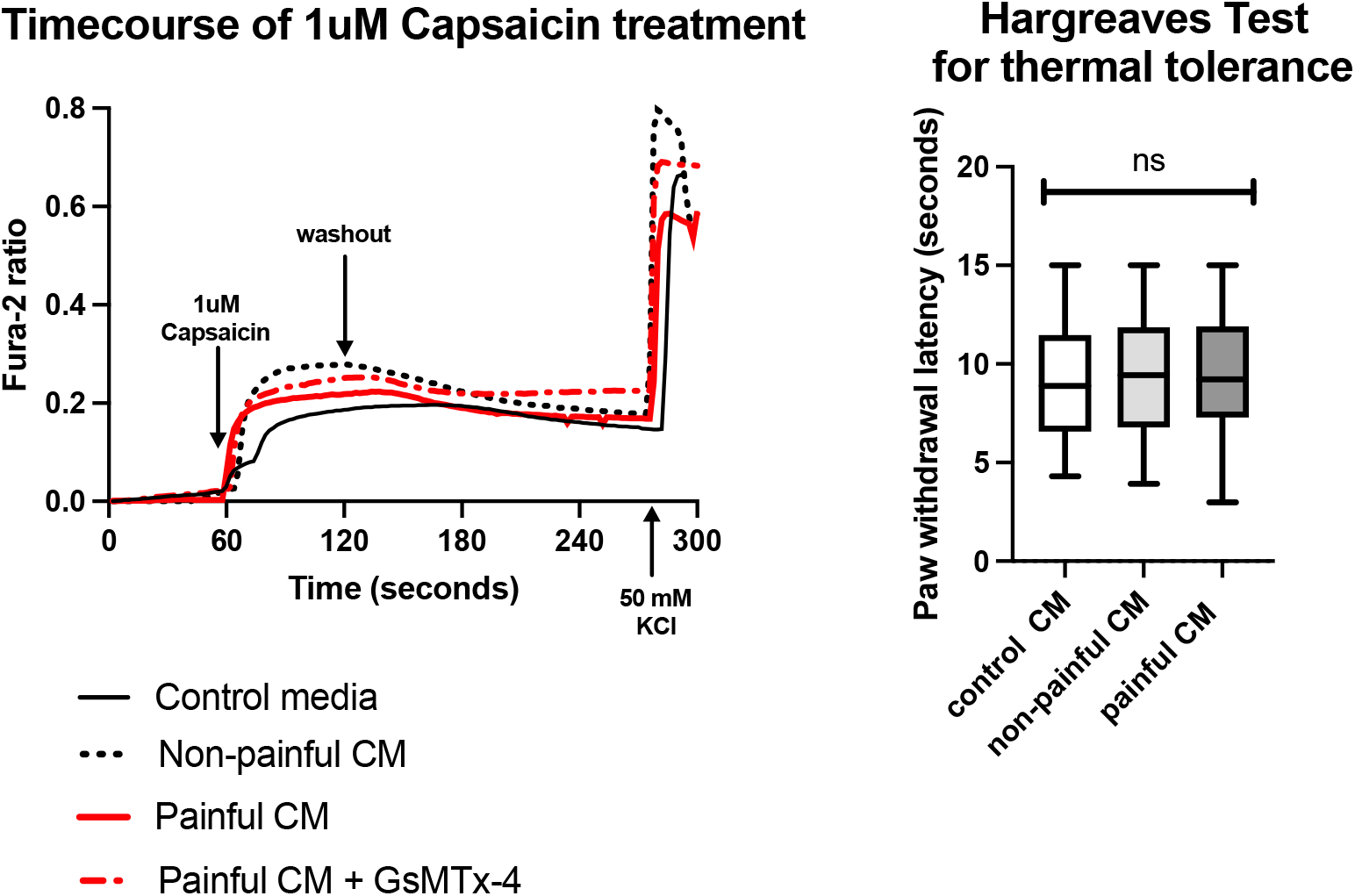
Painful conditioned media does not affect TRPV1 currents or thermal tolerance. **a)** Calcium imaging of CM treated DRG neurons Dissociated DRG cultures were pre-treated with painful schwannoma CM (+/-5uM GsMtx-4), non-painful schwannoma CM, or control media for 48 hours before imaging. Cells were perfused with the TRPV1 agonist Capsaicin (1uM) for 1 min with washout for 2 minutes prior to 50mM KCl. Fura-2 ratio measurements were recorded at 2 second intervals. The datapoints were corrected for differing baseline readings of Fura-2. Area under the curve was calculated for each treatment. Two coverslips of DRGs were tested per condition. No Differences were seen between treatment groups. *Brown-Forsythe ANOVA test* with Dunnett’s T3 multiple comparisons was employed to determine statistical significance (p<0.05 is considered significant) **b)** Hargreaves test for thermal tolerance Ten microliters of control media (n=10 mice) media from non-painful schwannoma (n=10 mice) and painful CM (n=10 mice) were injected into the footpad of C57Black mice. Paw withdrawal latency is recorded as the time it takes from the start of the heat stimulus for the mouse to lift its paw. No differences in paw withdrawal latency were detected between groups.

### Painful CM induces mechanically evoked pain in C57Black mice

We tested the effects of conditioned media on evoked pain by injecting the footpad of wild-type healthy C57Black mice and assessing for subsequent pain behaviors. Ten mice were injected with CM from painful tumors, 10 mice with media from non-painful (np) tumors, and 10 mice with non-conditioned (Control) medium (Dulbecco’s modified eagle’s medium/10% FBS/2uM forskolin). Samples were blinded to eliminate observer bias. The CM samples are indistinguishable in color, viscosity, and volume. Mice received two injections of media over the course of 48 hours. One hour after the final injection, the mice were examined for sensitivity to thermal sensitivity using the Hargreaves test **(Figure 3b)** or mechanically evoked pain using Von Frey filaments **(Figure 4)**. Painful schwannoma CM does not influence sensitivity to heat stimuli, as there were no differences in thermal tolerance observed among mice treated with control media, painful CM, or non-painful CM **(Figure 3b)**. These findings align with our in vitro calcium imaging results, which showed no differences in capsaicin sensitivity across cells treated with painful CM, non-painful CM, or control media **(Figure 3a)**.

**Figure 4:**
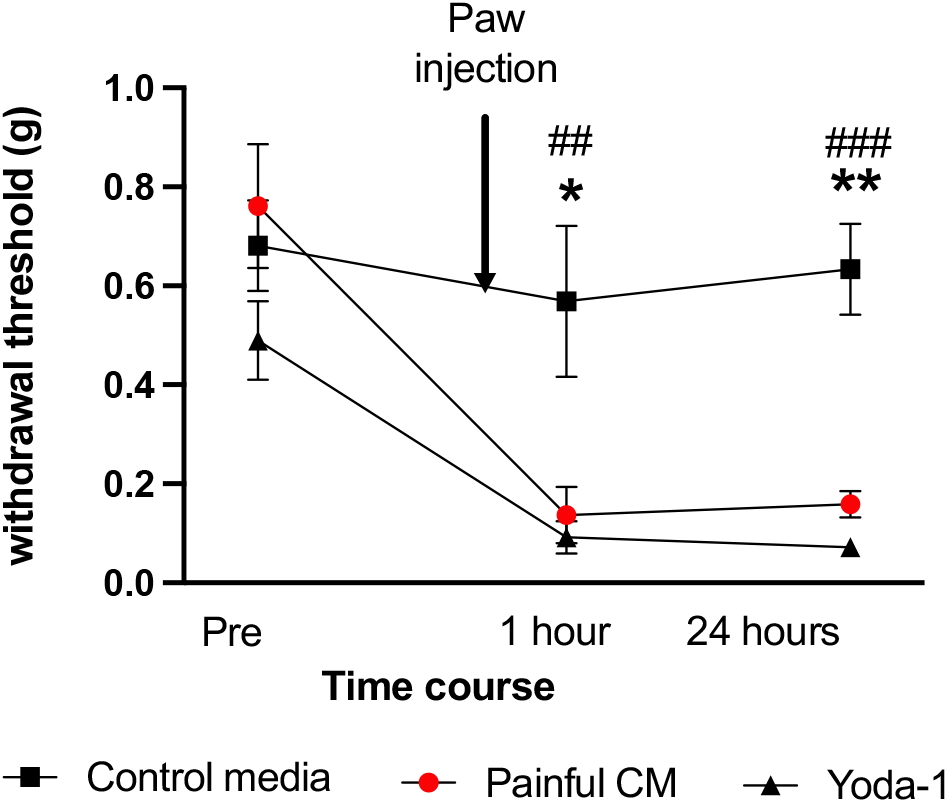
Yoda-1 and Painful CM caused a lasting hypersensitivity to mechanical stimuli. We induced mechanical pain via injection of 5uM Yoda-1 (an agonist of Piezo-1) into the footpad of C57Black mice. Yoda-1 was mixed with control media (Dulbecco’s modified eagle’s medium/10% FBS/2uM forskolin). The Von Frey simplified up/down method was used to assess mechanosensitivity. One hour post-injection of either painful CM (red circles) or control media +Yoda-1 (black triangles), mice demonstrated an increased response to mechanical stimuli as shown by a decrease in paw withdrawal threshold. (Pre-injection vs 1 hour post injection p=0.0001 painful CM; p=0.008 Yoda-1). Withdrawal threshold remained low for 24 hours post injection. Paw withdrawal threshold for mice injected with control media stayed constant over time. (## control vs Yoda-1 p=0.002 1 hour, ### control vs Yoda-1 p=0.0002 24 hours) (* control vs painful CM p=0.006 1 hour, ** control vs painful CM p=0.00124 hours). Two-away ANOVA with multiple comparisons was used to determine significance.

We examined the effects of painful CM and Yoda-1 on mechanical allodynia in our mouse model. Specifically, control media was combined with Yoda-1 at a final concentration of 5 μM. Ten microliters of painful CM, control media, or control media + Yoda-1 was injected into the footpad of one paw of a C57Black6 mouse. The simplified up/down (SUDO) method was used to assess hypersensitivity to light touch in the injected paw, utilizing Von Frey filaments ranging from 0.02 g to 1.4 g. This method determines the minimal force that consistently elicits a response, identifying the mechanical pain threshold.

Testing started with a mid-range force of 0.6 g. The filament was applied to the glabrous skin for 2 seconds, and a positive response was recorded if the mouse flinched, indicating a reaction to the stimulus. If a positive response occurred at 0.6 g, the next lower force was tested until no response was observed. Conversely, if the response was negative at 0.6 g, the next higher force was applied. Paw withdrawal threshold (PWT) was calculated using the simple up/down reader software [32], with pre-injection PWT averaging approximately 0.6g **(Figure 4)**.

Consistent with our previous findings [33], painful CM significantly enhanced mechanical stimulus-evoked behavioral responses compared to control media, as shown by a reduced paw withdrawal threshold one hour after injection **(Figure 4)**. The contralateral (non-injected) paw served as an internal control (data not shown). This heightened sensitivity persisted for 24 hours post-injection. Similarly, injecting 5 μM Yoda-1 into the footpad caused a comparable reduction in paw withdrawal threshold as painful CM, suggesting that CM-induced mechanotransduction is mediated by Piezo-1. This sensitization was sustained for 24 hours after injection **(Figure 4)**. In contrast, no changes in withdrawal thresholds were observed in mice treated with control media.

### GsMTx-4 prevents CM-induced mechanical hypersensitivity

As our data points to Piezo-1 involvement in SWN CM induced mechanosensitivity, we next tested the effect of GsMTx-4 on mechanical allodynia. We injected 10 uM GsMtx-4 into one footpad of C57Black mice mixed with control media injection (n=10 mice), control media+ 5uM Yoda-1 (n=10 mice) and Painful SWN CM (n=10 mice). One hour post-injection, Von Frey filaments were used to determine paw withdrawal threshold as previously described. GsMTx-4 prevented Yoda-1 induced mechanical hypersensitivity (gray dotted bar) as withdrawal threshold remained the same as uninjected mice **(Figure 5a)**. In our test group (n=10 mice) 10 uM GsMTx-4 mixed with painful SWN CM (red dotted bar) also prevented mechanical hypersensitivity in the Von Frey assay **(Figure 5a)**.

**Figure 5:**
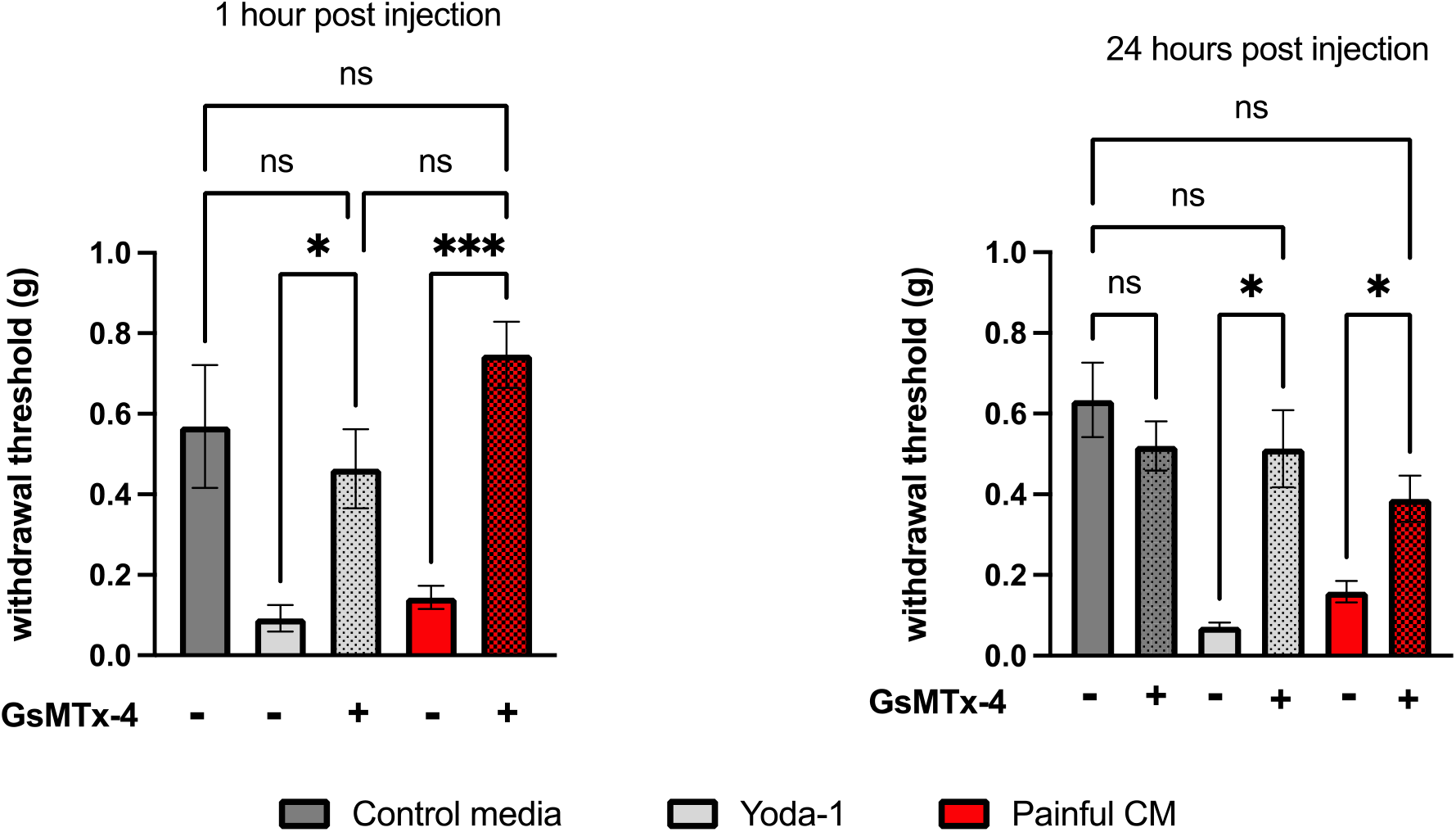
GsMtx-4 prevents and reverses sensitivity to mechanical force. **a)**. Prevention: the footpad of C57Black mice were injected with 10 ul of control media (n=10 mice), Yoda-1 +/-10 uM GsMTx-4 (n=10 mice), or Painful CM +/-10 uM GsMTx-4 (n=10 mice). One hour post injection mice were subjected to Von Frey testing. Mice receiving Yoda-1 or Painful CM showed a significant drop in withdrawal threshold. Mice receiving GsMTx-4 did not demonstrate a decrease in paw withdrawal level and were comparable to control media. **b)** Reversal: the footpad of C57Black mice were injected with control media (n=10 mice), Yoda-1 (n=10 mice), or Painful CM (n=10 mice). Twenty-four hours post injection mice were tested for mechanosensitivity using Von Frey filaments. Immediately following the VF, mice were injected with a ten microliter volume of 10uM GsMtx-4 (in control media). Thirty minutes post injection, Von Frey testing was repeated. Mice receiving GsMTx-4 showed a reversal in paw withdrawal threshold (Yoda-1 p=0.01, Painful CM p=0.01), demonstrating a rescue from the painful phenotype. GsMTx-4 had no effect on mice treated with control media.

### GsMTx-4 reverses CM-induced mechanical hypersensitivity

Painful CM or Yoda-1 caused a decrease in paw withdrawal threshold that is sustained for 24 hours as demonstrated by Von Frey testing **(Figures 4 &5b)**. Upon completion of the VF test this set of mice were next injected with 10uM GsMTx-4 and allowed to recover for 30 minutes. After 30 minutes another round of Von Frey testing was performed. Mice injected with GsMTx-4 demonstrated a significant reversal in paw withdrawal threshold in painful SWN CM or Yoda-1 groups **(Figure 5b)**. GsMTx-4 was not found to alter mechanical sensitivity of mice injected with control media, suggesting that activation of Piezo-1 is the manner in which painful CM induces mechanosensitivity.

## Discussion

Chronic pain is a significant issue for patients with schwannomatosis (SWN), profoundly impacting their quality of life [6]. Neuropathic pain medications often prove ineffective, while opioids merely mask the pain and carry substantial risks of adverse effects [6]. Despite a large tumor burden, not all tumors in a SWN patient cause pain[5]. Surgical removal of painful tumors is the standard treatment; however, this approach is not sustainable due to the potential need for multiple surgeries, along with the risks of nerve damage and other surgical complications [34]. Therefore, our focus is on developing safe, effective, and non-surgical therapies to manage pain in these patients.

One challenge in SWN research is the absence of an animal model capable of replicating tumor growth along peripheral nerves. To address this, we developed a surrogate pain model using healthy C57Black6 mice. Mice exposed to conditioned media (CM) from painful schwannoma tumors exhibited heightened sensitivity to mechanical stimuli [33]. Our lab investigates the interaction between tumor-secreted factors and sensory neuron pain pathways to identify targeted therapies for SWN [31]. While we have had success in detecting differences in gene expression in neurons exposed to tumor CM, notably, our findings on increased mechanosensitivity in CM-treated mice have led us to explore novel approaches to pain relief, including the use of GsMTx-4, an MSC inhibitor. Our in vitro studies demonstrated that GsMTx-4 effectively blocks Piezo1 currents induced by Yoda1 in dorsal root ganglion (DRG) cultures treated with painful CM **(Figure 2)**. Interestingly, CM exposure did not alter TRPV1 currents in vitro or affect thermal sensitivity in vivo **(Figure 3)**. Thus, targeting Piezo1-mediated mechanosensitivity with GsMTx-4 presents a logical therapeutic approach for SWN-related pain.

While we cannot directly model tumor growth on nerves, our system is functionally relevant. Piezo-1 is expressed in keratinocytes [35], sensory lanceolate endings [36], afferent axons and dorsal root ganglia **(Figure 1)** [37]. Activation of Piezo1 channels is associated with axon injury and inflammation [38]. A publication by Park et al 2008, showed that GsMTx-4 reduced mechanical allodynia that was induced by inflammation [39].

We have found elevated levels of proinflammatory cytokines and chemokines in CM from painful schwannomas [31, 33]. Therefore, we are confident that our assay models a tumor-induced inflammatory state that promotes mechanical allodynia. Interestingly, the Park et al study found that GsMTx-4 administration did not affect withdrawal thresholds to Von Frey filament stimulation when the tissue was not inflamed [39]. Our testing of GsMTx-4 in vivo suggests the same. GsMTx-4 had no effect in mice treated with non-conditioned media **(Figure 5)**. This is particularly promising as we aim to develop GsMTx-4 for treating inflammation related mechanical pain in SWN while minimizing off-target effects.

While GsMTx-4 significantly reduced pain sensitivity in our model, complete reversal of the painful phenotype was not achieved **(Figure 5b)**. Our experiments utilized a 10 µM concentration of GsMTx-4, whereas other in vivo studies employed concentrations ranging from 5 to 40 µM [15, 28, 37, 40-43]. We believe increasing the concentration or exploring the D-form of G
sMTx-4, which may be more potent, could achieve complete pain relief in our model. Future studies will focus on optimizing the dose-response and evaluating the efficacy of the D-form.

Our overall goal is to develop an effective treatment for SWN related pain that reduces the need for surgery. Local administration of GsMTx-4 near a tumor may be a minimally invasive and quickly translatable treatment for schwannomatosis related pain **(Figure 6)**. Peripheral Piezo1 and other mechanosensitive ion channels can be inhibited by sciatic nerve injection of GsMTx-4 [37]. Moreover, intradermal and intraperitoneal applications of GsMTx-4 have demonstrated anti-nociceptive effects against mechanical stimuli [39, 44] supporting its use to treat individual tumors that are causing pain.

**Figure 6:**
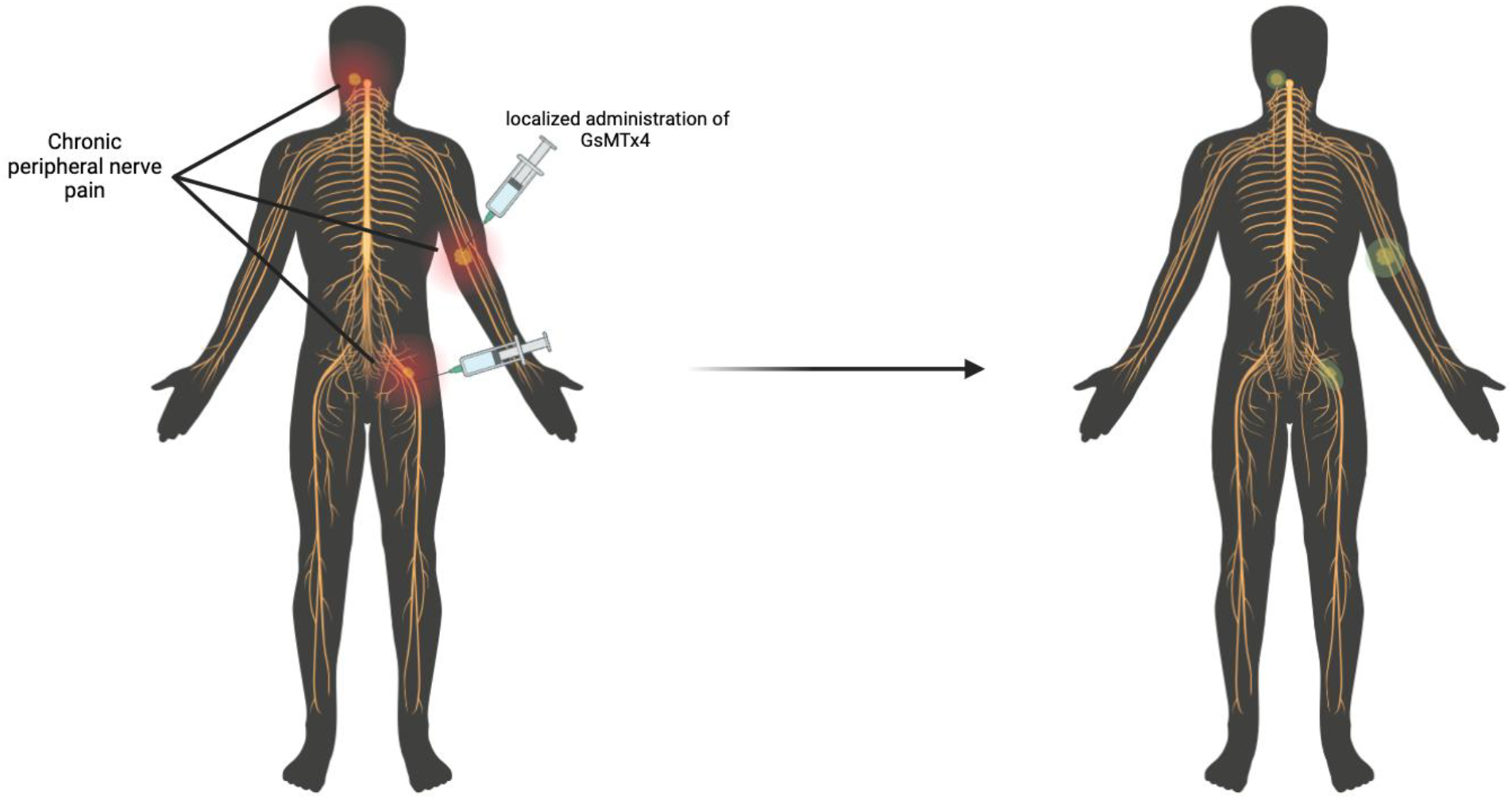
Patients with schwannomatosis develop multiple painful tumors along the spine and major peripheral nerves. Local administration of GsMTx-4 may be an effective minimally invasive treatment for tumor related pain.

The mechanism of action of GsMTx-4 and its D-form has been extensively studied in vitro and in vivo, demonstrating its effectiveness as an inhibitor of mechanosensitive channels in various disease contexts. [22, 23, 26, 28, 41]. Our current work, while still in its early stages shows that GsMTx-4 holds great promise as a useful alternative to surgical intervention for the treatment of SWN related pain.

## Supporting information

Supplementary stats

## Appendix

Statistical analysis was performed using Graphpad Prism 10. Pre-Hoc and Post-hoc testing results are included as an excel file.

## Author Contributions

C.G. conducted experiments, performed data analysis, performed troubleshooting to assist KLO in experimental design and co-wrote the manuscript. R.R. conducted experiments and performed data analysis, K.L.O. conceived the project, designed the experiments, conducted experiments, analyzed the data, and co-wrote the manuscript.

## Acknowledgements

The Children’s tumor Foundation, The Blaustein Pain Foundation, and The Johns Hopkins Neurosurgery Pain Research Institute supported this work. We thank members of the Caterina lab for helpful discussions.

## Data Availability

The authors will make materials, data and protocols available upon request.

Please contact the corresponding author, Kimberly L Ostrow PhD

